# Pupillometry and the vigilance decrement: Task-evoked but not baseline pupil measures reflect declining performance in visual vigilance tasks

**DOI:** 10.1101/2021.12.01.470724

**Authors:** Joel T. Martin, Annalise H. Whittaker, Stephen J. Johnston

**Author notes:** Author Note: Joel T. Martin, Department of Psychology, University of York, United Kingdom, YO10 5DD. The research was funded by a grant from the Dstl.

## Abstract

Baseline and task-evoked pupil measures are known to reflect the activity of the nervous system’s central arousal mechanisms. With the increasing availability, affordability and flexibility of video-based eye tracking hardware, these measures may one day find practical application in real-time biobehavioral monitoring systems to assess performance or fitness for duty in tasks requiring vigilant attention. But real-world vigilance tasks are predominantly visual in their nature and most research in this area has taken place in the auditory domain. Here we explore the relationship between pupil size—both baseline and task-evoked—and behavioral performance measures in two novel vigilance tasks requiring visual target detection: 1) a traditional vigilance task involving prolonged, continuous, and uninterrupted performance (*n* = 28), and 2) a psychomotor vigilance task (*n* = 25). In both tasks, behavioral performance and task-evoked pupil responses declined as time spent on task increased, corroborating previous reports in the literature of a vigilance decrement with a corresponding reduction in task-evoked pupil measures. Also in line with previous findings, baseline pupil size did not show a consistent relationship with performance measures. We discuss our findings considering the adaptive gain theory of locus coeruleus function and question the validity of the assumption that baseline (prestimulus) pupil size and task-evoked (poststimulus) pupil measures correspond to the tonic and phasic firing modes of the LC.

## 1 Introduction

The term *vigilance* has received varied usage in scientific research but broadly speaking it refers to an organism’s ability to sustain its attention over prolonged periods of time (Kahneman & Treisman, 1984; Raja Parasuraman et al., 1998; Raja Parasuraman & Davies, 1982; Joel S. Warm et al., 2008; Joel S. Warm & Jerison, 1984). Although there is a long history of research into performance and continuous work tasks (see Bills, 1943; Hockey, 2013 for reviews), Mackworth (1948, 1950) is frequently credited for the first systematic studies of vigilance and the discovery that human detection performance on a monotonous watch keeping task, under conditions similar to those experienced by radar and sonar operators, declines as time spent on-task increases. This so-called *vigilance decrement* became the target of numerous research efforts in human factors and experimental psychology which sought to understand how factors specific to the task, the individual performing it, and the environment in which it is performed, all contribute to failures of vigilant attention (Frankmann & Adams, 1962; Mackie, 1987; Wiener, 1987). Signal detection theory (Green & Swets, 1974) has played a central role in the psychophysical analysis of vigilance studies, with detection performance being characterized frequently on the basis of the number of hits, misses, false alarms, correct rejections, and the derived measures of sensitivity and criterion (e.g., Mackworth, 1970; Parasuraman & Davies, 1976). Detection latency also features in analyses of vigilance task performance (e.g., Basner & Dinges, 2011; Broadbent, 1958; Buck, 1966) and biometric technologies such as electroencephalography (EEG) and functional magnetic resonance imaging (fMRI) continue to shape our understanding of the neurophysiological mechanisms of vigilant attention (for review, see: Fortenbaugh, DeGutis, & Esterman, 2017; Langner & Eickhoff, 2013; Oken, Salinsky, & Elsas, 2006).

Another biometric technique that has been successfully applied to the study of vigilant attention is cognitive pupillometry, the measurement of the size and reactivity of the eyes’ pupils following exposure to psychologically relevant stimuli. The pupils respond primarily to light, but when light levels are held constant, fluctuations in pupil size offer a window of insight into the brain’s central arousal systems (Joshi & Gold, 2020; Kahneman, 1973; Laeng et al., 2012). Specifically, nonluminance-mediated pupil size changes are known to reflect the moment-to-moment activity of the locus coeruleus noradrenalin system (LC-NA: Joshi et al., 2016; Rajkowski, Kubiak, & Aston-Jones, 1993), which has a central role in the modulation of arousal and alertness (Berridge, 2008; Berridge et al., 2012; Berridge & Waterhouse, 2003) and in maintaining optimal levels of vigilance and performance (Petersen & Posner, 2012; Posner & Petersen, 1990). Extensive single-cell recording studies in behaving rodents and monkeys show that the noradrenergic neurons of the LC exhibit *phasic* and *tonic* modes of activation and that these distinct modes correspond to different behavioral states (Aston-Jones & Cohen, 2005; Aston-Jones, Rajkowski, & Cohen, 1999). The phasic mode is characterized by short bursts of activation in response to task-relevant stimuli and supports task engagement and exploitation of the environment, whereas the tonic mode is characterized by a sustained increase in baseline activation in response to diminishing task utility and supports disengagement from the current task and exploration of the environment (Aston-Jones & Cohen, 2005).

The functional role of the LC-NA system and its association with the pupil has led many to assume that baseline (i.e., prestimlus) and task-evoked measurements of pupil size may correspond to the tonic and phasic firing modes of the LC, and that these measures may reflect changes in vigilant attention over time. Indeed, changes in pupillometric response associated with changes in vigilant attention have been noted previously. Beatty (1982) asked participants to monitor a string of tones presented at 3.2 s intervals continuously for 48 min for the occurrence of target tones, which were slightly attenuated in volume. Approximately 12 targets were presented at random intervals in every 5 min period, with 108 targets being presented across the whole task. As is common in vigilance tasks, while the task itself was conducted as one continuous 48-minute procedure, the data were sub-divided into several periods of watch and compared to determine time-related differences in performance. Detection accuracy decreased in accordance with time spent on task, replicating the classic finding of a vigilance decrement described by Mackworth (1948, 1950). Task-evoked pupillary responses to target stimuli mirrored these results, decreasing in amplitude across each third of the test; but baseline (prestimulus) measurements of pupillary activity (obtained prior to each target stimulus) showed little change. In more recent work we even see hints that pupil measures may serve to predict performance on a moment-to-moment basis. For example, Kristjansson et al. (2009) reported significant differences in pupil size and dilation rate for the fastest and slowest detection responses in a psychomotor vigilance task, suggesting that the pupil measures may provide sufficient reliable information to index alertness in real-time. But, despite the promising narratives of Kristjansson et al. and a handful of other studies (e.g., Unsworth & Robison, 2016; van den Brink et al., 2016), the nature of the relationship between pupil and performance measures remains unclear.

Since Beatty (1982), many studies have found performance decrements in long and demanding tasks that coincided with reduced task-evoked pupil responses (e.g., Hopstaken, van der Linden, et al., 2015; Hopstaken, Van der Linden, et al., 2015; Murphy, van Moort, & Nieuwenhuis, 2016; Unsworth & Robison, 2016) but, as noted by van den Brink et al. (2016), the literature is conflicted on the relationship between task performance and baseline pupil size. In some experiments, moments of off-task thought or poor task performance were associated with larger pupils at baseline (Franklin et al., 2013; Gilzenrat et al., 2010; Smallwood et al., 2011, 2012; Unsworth & Robison, 2016), whereas in other experiments poor task performance was associated with smaller pupils at baseline (Grandchamp, Braboszcz, & Delorme, 2014; Hopstaken, Van der Linden, et al., 2015; Kristjansson et al., 2009; Mittner et al., 2014; Van Orden, Jung, & Makeig 2000), or occurred following a gradual decrease in baseline pupil size (Grandchamp et al., 2014; Massar et al., 2016; McIntire, McKinley, & Goodyear, 2014; Murphy et al., 2011). Poor task performance was also found to occur with both relatively large and small baseline pupil size within experiments (van den Brink et al., 2016; Murphy et al., 2011; Smallwood et al., 2012; Unsworth & Robison, 2016), with one experiment reporting an increase in baseline pupil diameter as a function of time-on-task in a 37-min auditory vigilance task without breaks (Murphy et al., 2011).

Such discrepant findings on the relationship between baseline pupil size and task performance likely reflect the interplay of various methodological factors. Among the studies cited in the previous paragraph there is considerable variability in how performance was measured, with some focusing primarily on RT measures, such as mean RTs (e.g., Smallwood et al., 2011, 2012), fraction of the slowest or fastest RTs (e.g., Unsworth & Robison, 2016; van den Brink et al., 2016), or RT variability (e.g., Murphy et al., 2011); and others focusing more on perceptual sensitivity (i.e., d*’* : Beatty, 1982; Hopstaken, van der Linden, et al., 2015; Hopstaken, Van der Linden, et al., 2015), or self-reported measures of task engagement (e.g., Franklin et al., 2013; Grandchamp et al., 2014; Mittner et al., 2014). Task demands also vary considerably across experiments, with some requiring only simple target detection (e.g., Massar et al., 2016) and others requiring *simultaneous* (Beatty, 1982; van den Brink et al., 2016; Gilzenrat et al., 2010; Murphy et al., 2011) or *successive* (Hopstaken et al., 2016; Hopstaken, van der Linden, et al., 2015; Hopstaken, Van der Linden, et al., 2015; Smallwood et al., 2011, 2012) discrimination^1^. Further, some tasks called for prolonged continuous monitoring (Beatty, 1982; Murphy et al., 2011), whereas others entailed intermittent breaks from the primary task (e.g., Hopstaken, Van der Linden, et al., 2015; Smallwood et al., 2004; Unsworth & Robison, 2016), which even when very short have the potential to improve performance by temporarily boosting motivation (Ariga & Lleras, 2011; Ralph et al., 2016; Ross, Russell, & Helton, 2014). Finally, the stimuli varied substantially, and some may have had undesirable behavioral or pupillometric consequences. For example, the ‘running counter’ stimulus used in Massar et al.’s (2016) psychomotor vigilance task provides feedback which could enable participants to detect declines in their performance and adopt compensatory strategies (Thorne et al., 2005); and for studies using visual stimuli (e.g., van den Brink et al., 2016; Smallwood et al., 2011, 2012), differences in visual attributes such as luminance, color and contrast may have contributed to pupillometric and behavioral variance (Barbur, Harlow, & Sahraie, 1992; Goldwater, 1972).

The increasing availability, affordability, and flexibility of video-based eye tracking hardware means that pupils’ predictive power for vigilant attention may one day find practical application in passive, real-time biobehavioral monitoring systems to assess performance or fitness for duty. Such systems would be of particular use in scenarios where traditional reaction time assessments cannot easily be administered. Presently, however, it remains unclear which pupil measures, if any, would be suitable for an application of this kind. Focusing on the issues raised above, here we examine the relationship between pupil measures—both baseline and task-evoked—and task performance in a novel implementation of two well-established vigilance task paradigms: 1) continuous, uninterrupted vigilance, and 2) psychomotor vigilance.

## 2 Experiment 1

Since Mackworth (1948, 1950), experimental vigilance tasks have generally aimed to simulate the conditions of real-world scenarios where monotonous repetitive tasks have become commonplace due to automation and industrial mechanisation. Though vigilance tasks can vary in many ways, the defining characteristic is that observers must remain alert and respond to critical signals presented against a background of noncritical signals over prolonged, unbroken stretches of time—usually at least 30 mins (Frankmann & Adams, 1962; Parasuraman & Davies, 1976). Key differences between tasks known to influence performance are the sensory modality of stimulus presentation (e.g., auditory, visual), the psychophysical dimensions used to define critical signals (e.g., brightness, loudness), and whether the detection of targets requires successive or simultaneous discrimination (Parasuraman, 1979; Warm et al., 2008); but performance ultimately depends on complex interactions between factors relating to the task, the environment, and the individual (Ballard, 1996). To date, most vigilance tasks conducted with pupillometry have presented stimuli in the auditory modality, probably to avoid methodological confounds associated with the effects of visual stimulation and optical distortion of raw pupil measurements inherent to video-based systems. But vigilance tasks in the real word are predominantly visual, and if pupil measures are to serve a useful purpose for tracking vigilant attention in real world settings, they must be robust to the effects of visual stimuli.

In Experiment 1 we explored the relationship between pupil and performance measures (RT, accuracy, *d’* and *c*) in a canonical vigilance task with stimuli presented in the visual modality. To our knowledge, McIntire et al. (2014) is the only previous example of such a study, though the analysis was correlational and simply explored how average pupil size and percentage of hits showed a similar decline across four successive 10-min periods of continuous task performance. This is in contrast to auditory vigilance experiments (e.g., Beatty, 1982; Gilzenrat et al., 2010; Murphy et al., 2011), where there has been considerable focus on event-related pupil measurements. The present experiment therefore aimed to examine event-related pupil responses in a novel vigilance task with visual stimuli, whilst controlling appropriately for the effects of eye movements and luminance confounds.

Our task required participants to continuously monitor four centrally presented equiluminant visual stimuli for 30 min in order to detect and respond to brief targets occurring with temporal and spatial uncertainty against a high background event rate. A relatively high number of targets—6 per min—was used to ensure that a suitable amount of event-related data would be generated for the analysis (e.g., Mackie, 1987). We predicted that performance measures (e.g., RT, accuracy, *d’*) across successive 10 min task blocks would betray a classic vigilance decrement as has been reported widely in the literature (Frankmann & Adams, 1962; Mackie, 1987; Mackworth 1948, 1950; Wiener, 1987). Second, based on the most consistent findings from pupillometric studies of vigilance (e.g., Hopstaken, van der Linden, et al., 2015; Hopstaken, Van der Linden, et al., 2015; Murphy, van Moort, & Nieuwenhuis, 2016; Unsworth & Robison, 2016), we predicted that the magnitude of task-evoked pupil size changes would decrease, in line with performance measures, across the duration of the task. Considering the discrepant findings in the literature, we did not make specific predictions about baseline or ‘tonic’ pupil size for this experiment.

### 2.1 Materials and methods

#### 2.1.1 Participants

Twenty-eight participants (23 females; age range 18-32 years, *M* = 20.07, *SD* = 2.8) completed the experiment voluntarily or in exchange for course credit. All participants were students at Swansea University reporting normal or corrected-to-normal acuity and color vision. The experimental protocol was approved by the Ministry of Defence Research Ethics Committee and the Department of Psychology Ethics Committee at Swansea University. Written informed consent was obtained from each participant.

#### 2.1.2 Design

Task design reflected the core principles of classic experimental vigilance paradigms (Baddeley & Colquhoun, 1969; Mackworth, 1948, 1950; Parasuraman & Davies, 1976). Participants were asked to monitor four low-contrast gratings arranged squarely around a central fixation circle. The gratings rotated synchronously in a clockwise ticking motion at a rate of 120 ticks per minute (30° rotation per tick) and targets were defined as instances where one became briefly out of sync with the others (i.e., it missed a tick: see Figure 1). The task lasted for 30 min, during which time continuous monitoring was required. Six targets were presented every minute (180 overall) at pseudorandom intervals, subject to the following constraints: 1) the time between targets was at least 6 s and at most 30 s, 2) targets did not occur within 2 s of the beginning or end of the task. Targets occurred equally often at all of the four locations, although this was randomized across the whole experiment so that spatial uncertainty as to the location of the target would contribute to task difficulty (Broadbent, 1958; Mackie, 1987; Warm, Parasuraman, & Matthews, 2008). All participants completed one trial of this experiment in a single testing session lasting approximately 40 min.

**Figure 1.**
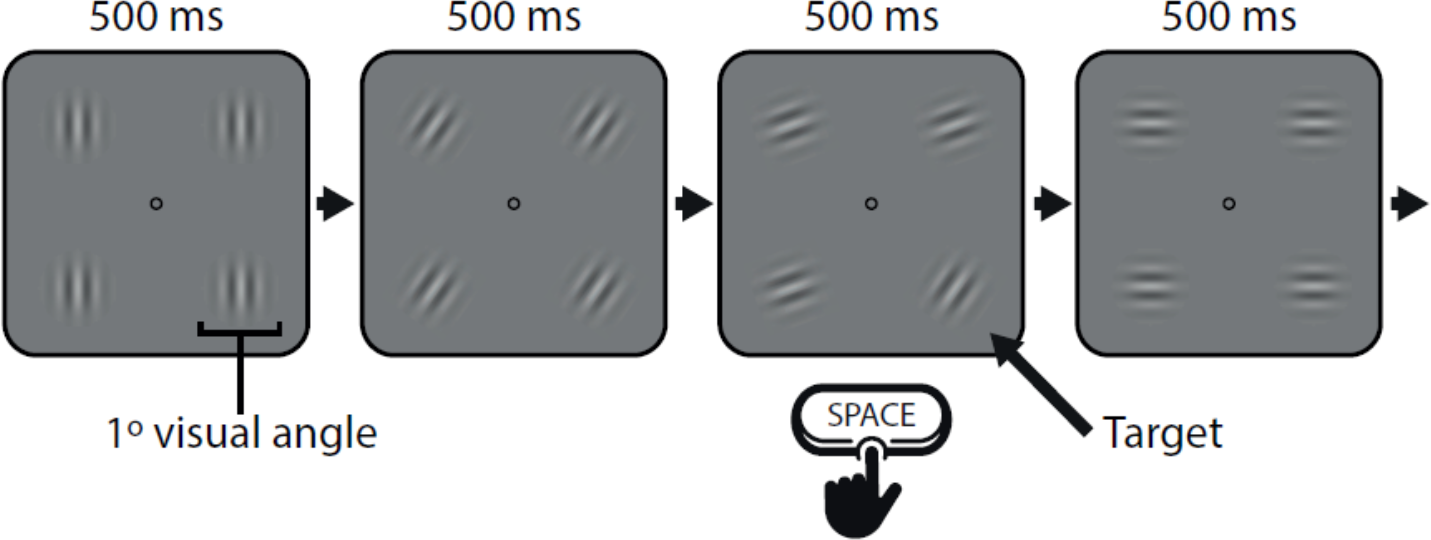
Stimuli and trial sequence for Experiment 1. Participants were asked to respond by pressing space every time one of the gratings did not follow the standard sequence (6 times per min).

#### 2.1.3 Stimuli and apparatus

Four gratings enveloped within a cosine window (spatial frequency = 0.1, *SD* = 12, 39% contrast), each spanning 1° × 1° of visual angle, were arranged in a square pattern around a central fixation circle on a grey background (Figure 1). The gratings were generated using an online tool^2^ and saved in JPEG format. The task was administered on a 24-in. Ilyama monitor running at a resolution of 1024 × 768 (1:1 aspect ratio) with a refresh rate of 144 Hz, and button responses were collected on a standard computer keyboard. The monitor and eye tracker were enclosed such that the only direct illumination came from the display screen and the participant could not see anything in their periphery. A viewing distance of 40 cm was maintained by a chin rest and forehead bar. Using a colorimeter (ColorCAL MKII, Cambridge Research Systems), the surface luminance of the grey background was recorded as 73.54 cd/m^2^, and the surrounding dark light of the unused portion of screen as 0.53 cd/m^2^. Pupil size and gaze data were recorded monocularly (left eye) with an EyeLink 1000 (SR Research, Mississauga, Ontario, Canada) system in tower mount configuration (32 mm lens) sampling at 250 Hz. According to the user manual, the system resolves pupil diameter to within 0.2 mm (SR Research Ltd., 2010). Eye level and camera position remained constant throughout the recording session for each participant. Stimulus presentation was managed with Experiment Builder (SR Research, Mississauga, Ontario, Canada).

#### 2.1.4 Procedure

On arrival at the lab, participants were told that for the next 30 min they would be required to complete a vigilance task that involved monitoring four circular patches rotating with a ticking motion at the center of the screen. It was explained that, from time to time, one of the patches would briefly become out of phase with the others, and that this was a target to which they had to respond. Participants were not given any further information about the frequency or temporal and spatial uncertainty of the targets. Once comfortable with the definition of a target, they were instructed that their task was to press the space bar every time they noticed such an event. Participants were forewarned that the task was monotonous, but were asked to try and respond as quickly and accurately as possible. They were also instructed to maintain central fixation on the screen. A 5-point calibration and validation routine was performed prior to starting the experiment.

#### 2.1.5 Task performance and pupillometry

Task performance was assessed with RT, accuracy (i.e., percent hits and false alarms), and the signal detection theory measures sensitivity (*d’*) and response bias (*c*). Hits, correct rejections, misses and false alarms were determined by an iterative algorithm which assigned button responses to stimulus events. For each button response, the RT in milliseconds to the last target was calculated. If the RT was greater than the minimum time between targets (6 s), or if no target had yet been presented, the button response was allocated to the nearest elapsed neutral event and labelled as a false alarm. All remaining button responses were then grouped together and a permissible range for hits was determined as ± 2 median absolute deviations (Leys et al., 2013) from the group-level median RT. Accordingly, all button responses that occurred within 225 to 1156 ms of targets were counted as hits, and those with RTs outside this range, as previous, were allocated to the nearest neutral event and counted as false alarms. The resulting distribution of RTs and the permissible hit range is illustrated in Figure 2. Finally, targets without a valid button response were counted as misses, and all remaining neutral events as correct rejections. This process of dealing with behavioral responses in sustained-attention tasks with high event rates is similar to that used by Esterman et al. (2016) and van den Brink et al. (2016). For the purposes of analysis, the complete experiment was decomposed into three 10 min periods of watch.

**Figure 2.**
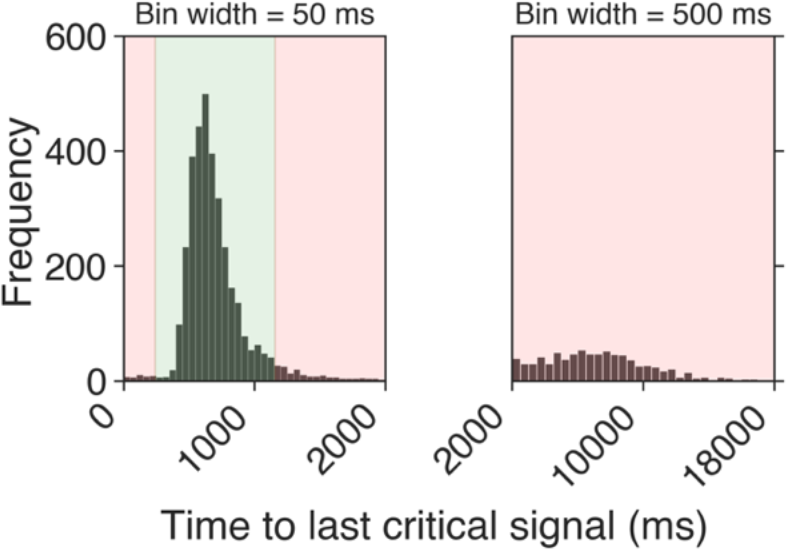
Distribution of RTs for Experiment 1 across all participants in 50 ms bins (left) and 500 ms bins (right). The green shaded area demarcates the permissible hit range (225 to 1156 ms after the target), and the red shaded areas show the range where button responses were considered false alarms.

Individual pupil traces were extracted for all signal detection theory outcomes. For misses and correct rejections the pupil data were time locked to the stimulus event and baseline pupil size was defined as the average pupil size in a 500 ms window prior to the event. For hits and false alarms, the data were time locked to the corresponding button event and the baseline period was offset by a further 500 ms (i.e., from -1000 to -500 ms prior to the button response) to minimize contamination from fluctuations in pupil size associated with movement preparation and execution (Einhäuser, Koch, & Carter, 2010; Hupé, Lamirel, & Lorenceau, 2009; Martin, Whittaker, & Johnston, 2020; Richer & Beatty, 1985).

Baseline and task-evoked pupil measures were also derived from the pupil traces to probe the effects of time-on-task. Baseline measures were defined as the average of the *z*-transform of pupil size (across the whole experiment) in the baseline periods, and task-evoked measures were defined as the average percent modulation in the peristimulus intervals (i.e., the portion of the pupil trace not including the baseline). We also averaged the *z*-transform of pupil size into 1-min bins for an overarching look at how pupil size varied across the whole task.

#### 2.1.6 Data processing and statistical analysis

Pupil data were processed and analyzed using custom python scripts. Eye-blinks were detected using the standard EyeLink parsing algorithm and reconstructed with linear interpolation prior to smoothing with a 3rd-order Butterworth filter (4 Hz cut-off). Frequently, pre- and post-blink samples were noticeably part of the blink artifact, so we extended the blink endpoints by 100 ms in each direction. The average amount of data replaced by blink interpolation across all participants that were included in the analysis was 6.5%. Pupil data were then down-sampled to 50 Hz, baseline corrected at the trial level with the subtractive procedure (Mathôt et al., 2018), and converted to units of percent signal change.

To examine the general pattern of pupil measures for each of the signal detection theory outcomes, stimulus- (misses and correct rejections) and button-locked (hits and false alarms) pupil traces from across the whole experiment were compared using two-tailed nonparametric permutation tests with cluster-based correction for the multiple comparisons problem (Maris & Oostenveld, 2007). This approach does not depend on theoretical assumptions about the data and reduces experimenter bias associated with choosing a time-period over which to compute summary statistics. For the permutation tests, *t*-tests were used to compare two conditions (significance thresholds for test statistics determined theoretically from the appropriate degrees of freedom at *a* = .05) and we follow the guidance of Sassenhagen and Draschkow (2019) for reporting and interpretation. To examine time-on-task effects, performance (RT, accuracy, *d’*, *c*) and scalar pupil measures were averaged within 10-min watch periods and analyzed with repeated measures ANOVAs. Where Mauchly’s *W* indicated that the assumption of sphericity was violated, *p*-values were adjusted using the Greenhouse-Geisser correction. An alpha level of .05 was used for all statistical tests. The mean and standard deviation of horizontal (*M* = 527, *SD* = 20) and vertical (*M* = 379, *SD* = 26) gaze position for all samples included in the analysis indicate that participants maintained steady fixation at the center of the screen throughout the task.

#### 2.1.7 Exclusions

Pupil data associated with stimulus and button events were discarded if the participant blinked during the baseline period or if more than 25% of the data across the epoch of interest were interpolated. Overall, this led to the discarding of pupil data for 17.89% of stimulus-locked (i.e., misses and correct rejections) epochs and 14.32% of button-locked (i.e., hits and false alarms) epochs. No participants were excluded from the analysis.

### 2.2 Results

#### 2.2.1 Task performance

The average number of button responses made during the task was 151 (*SD* = 51). With respect to target events, the average percentage of hits and false alarms was 63.77% (*SD* = 14.66%) and 1.09% (*SD* = 1.43%), respectively. These data, together with the remaining behavioral data, are summarized in Figures 2 and 3. Figure 2 shows the RT distribution for the whole experiment including the permissible hit range (225 – 1156 ms post-target), and Figure 3 shows accuracy (percentage of hits and false alarms), RT for hits, sensitivity (*d’*) and response bias (*c*) as a function of Watch Period. The average RT for all hits was 666 ms (*SD* = 156 ms). Average sensitivity and response bias across the whole experiment were 2.93 (*SD* = 0.67) and 1.07 (*SD* = 0.26), respectively, indicating that perceptual sensitivity to targets was good, but also that participants were generally biased to withhold responses to targets.

**Figure 3.**
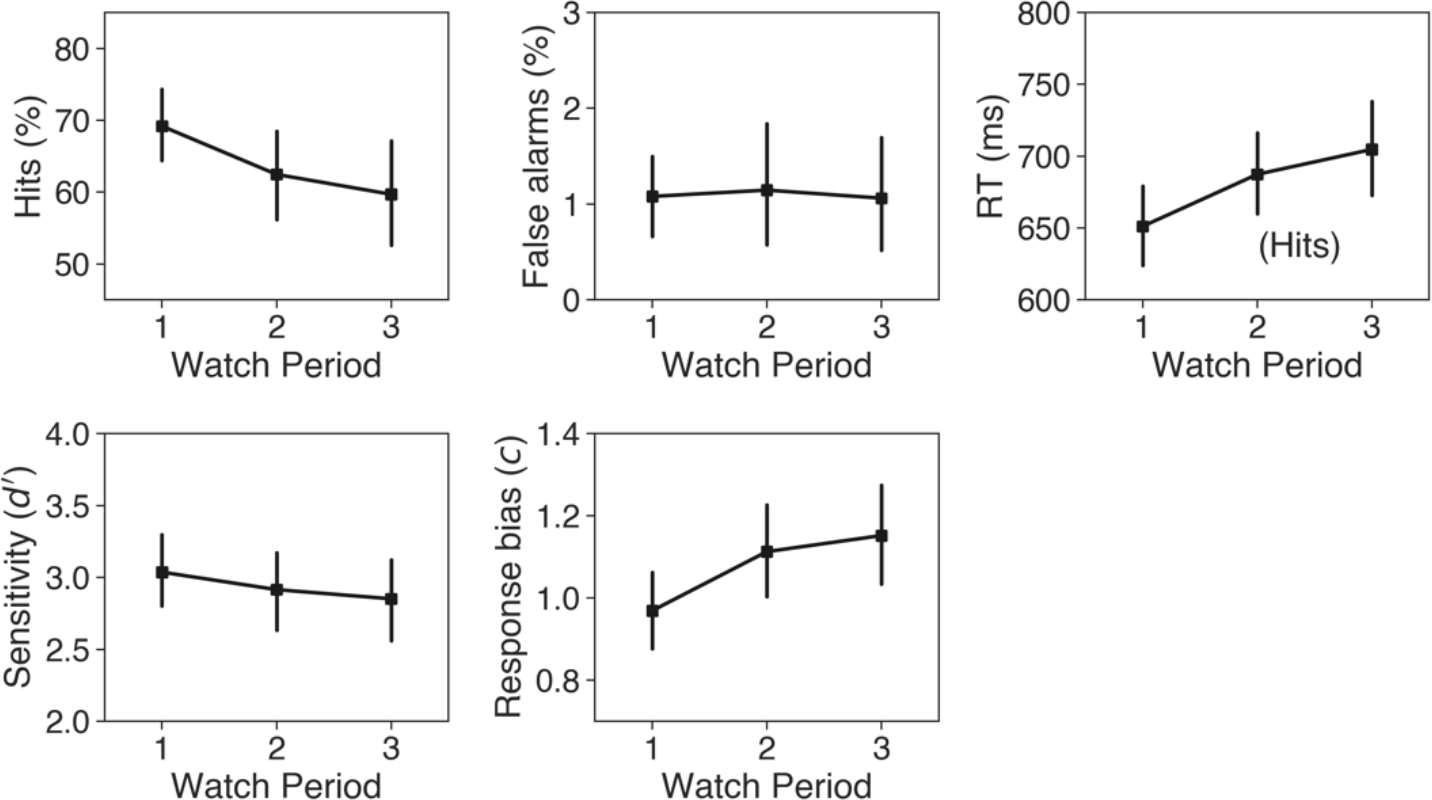
Task performance as a function of Watch Period in Experiment 1: percentage of hits (top-left) and false alarms (top-middle), RTs for hits (top-right), sensitivity (bottom-left) and response bias (bottom-middle). Error bars reflect 95% confidence intervals (bootstrapped, 1000 iterations).

To examine the effects of time-on-task, one-factor (Watch Period) repeated measures ANOVAs were conducted for all performance measures. First, the percentage of hits and false alarms were analyzed. There was a significant main effect of Watch Period on the percentage of hits, *F*(2, 54) = 6.14, *p* = .004, *ηp^2^* = 0.19, with Bonferroni-corrected *t*-tests showing that participants attained a significantly higher percentage of hits in Watch Period 1 (*M* = 69.2%, *SD* = 14.0%) compared to Watch Period 3 (*M* = 59.6%, *SD* = 19.7%), *t(28)* = 2.96, *SEM* = 3.21, *p* = .019. The difference between Watch Period 1 and Watch Period 2 (*M* = 62.46%, *SD* = 16.7%) was marginally significant, *t(28)* = 2.37, *SEM* = 2.83, *p* = .076, and the remaining comparison (2 vs. 3) was not significant (*p* = .662). The effect of Watch Period on the percentage of false alarms was not significant (*p* = .894).

ANOVA on the RT data for hits revealed a significant main effect of Watch Period, *F*(2, 54) = 16.51, *p* < .001, *ηp^2^* = 0.38. Post hoc analysis with Bonferroni adjustment confirmed that RT was significantly faster in the first Watch Period (*M* = 651 ms, *SD* = 79 ms) compared to the second (*M* = 687 ms, *SD* = 78 ms), *t(28)* = 4.95, *SEM* = 7.31, *p* < .001, and the third (*M* = 704 ms, *SD* = 89 ms), *t(28)* = 5.47, *SEM* = 9.79, *p* < .001, but that there was no significant difference in RT between Watch Period 2 and Watch Period 3 (*p* = .384).

The same analysis was repeated for the signal detection measures sensitivity (*d’*) and response bias (*c*). There was no significant main effect of Watch Period on sensitivity (*p* = .193), indicating that participants’ ability to discriminate targets did not change throughout the task, but there was a significant main effect on response bias, *F*(1.64, 44.31) = 7.38, *p* = .003, *ηp^2^* = 0.22. Bonferroni-corrected post hoc *t*- tests showed that response bias in Watch Period 1 (*M* = 0.96, *SD* = 0.26) was significantly lower than it was in Watch Period 2 (*M* = 1.11, *SD* = 0.31), *t*(28) = 2.8, *SEM* = 0.05, *p* = .028, and Watch Period 3 (*M* = 1.15, *SD* = 0.33), *t*(28) = 3.11, *SEM* = 0.06, *p* = .013, but that there was no significant difference between Watch Period 2 and Watch Period 3 (*p* = .937). This suggests that participants became more conservative as the task progressed and were therefore more reluctant to report that a target was present. As predicted, these performance data are consistent with the classic vigilance decrement.

#### 2.2.2 Pupil data

The *z*-transform of pupil data declined sharply in the first few minutes of the task and then increased steadily until the end. A cluster in the observed data extending from the 4^th^ to the 12^th^ minute differed significantly from the population mean (top panel, Figure 4). This time-on-task effect is well-noted in the literature for many different types of experiment (Fried et al., 2014; Hopstaken et al., 2016; Hopstaken, Van der Linden, et al., 2015; Massar et al., 2016; McIntire et al., 2014; Unsworth & Robison, 2016; van den Brink et al., 2016) and may reflect changes in overall arousal state or, more specifically, the transition from phasic to tonic modes of LC output (Joshi & Gold, 2020).

**Figure 4.**
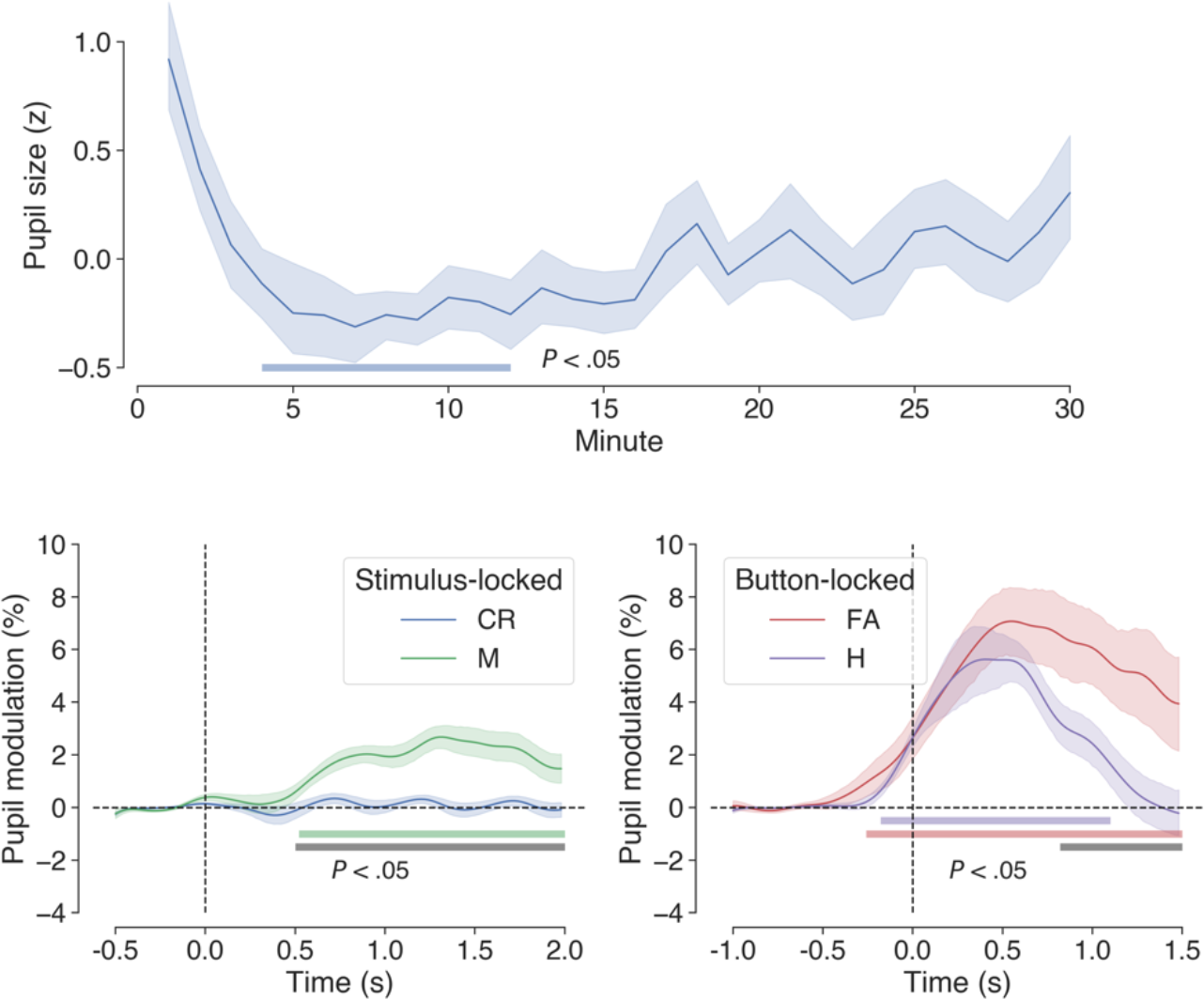
Grand average pupil data in Experiment 1. The top panel shows the average *z*-transformed pupil data between subjects in 1-min bins across the whole experiment. A cluster extending from the 4^th^ to the 12^th^ minute differed significantly from the population norm (blue colored bar). The bottom panels show stimulus- (left: correct rejections and misses) and button-locked (right: hits and false alarms) averages expressed as %-change from baseline, with colored horizontal bars indicating clusters of significant modulation from baseline and gray bars showing significant differences between traces (1024 permutations, *p* < .05, cluster-corrected for multiple comparisons). Shaded areas surrounding the pupil traces denote the standard error of the mean (SEM: bootstrapped, 5000 iterations).

Event-related pupil data were time-locked to button events for hits and false alarms and to stimulus events for misses and correct rejections. This was to ensure the comparability of pupil data that were consistently affected by motor acts. The grand-average pupil traces for each of these behavioral outcomes are shown in Figure 4. Both button-locked outcomes (hits and false alarms) showed the usual pattern of pupil modulation associated with the preparation and execution of motor responses (e.g., see Einhäuser et al., 2010; Hupé et al., 2009; Martin et al., 2020; Richer & Beatty, 1985), with dilation beginning up to 500 ms before the motor act and peaking shortly afterwards. Permutation tests revealed significant modulation from baseline for hits and false alarms, as well as a significant difference between these two outcomes (lower-right panel of Figure 4). The differences between the two traces can be summarized as follows. For hits, there was an average pupil modulation of 2.04% and a peak modulation of 5.62% with a latency of 400 ms from the button press, whereas for false alarms these values were 3.55%, 7.07%, and 540 ms, respectively. For the stimulus-locked pupil measures (misses and correct rejections), only misses resulted in significant modulation from baseline in the poststimulus period, and there was also a significant difference in pupil modulation between misses and correct rejections (bottom-left panel of Figure 4). For misses, there was an average modulation of 1.25% and a peak modulation of 2.68% with a latency 1320 ms. The time-course for correct rejections resembled a flattened sine wave (average modulation of -0.05%, peak modulation of 0.34%) in phase with the onset of stimulus events.

This periodic pattern is redolent of van den Brink et al.’s (2016) pupillometry data, which were observed in a task with a similar event-related design. We attribute this to task-correlated blinking and the blink-induced pupillary response (Knapen et al., 2016).

After examining the stimulus- and button-locked pupil traces across the whole experiment we probed the effects of time-on-task by analyzing the scalar representations of pupil data with two-way (Outcome × Watch Period) repeated measures ANOVA. These data are displayed in Figure 5. For the stimulus-locked (i.e., misses and correct rejections) baseline pupil data, there were no significant effects of Watch Period (*F*[2, 54] = 1.99, *p* = 0.146) or Outcome (*F*[1, 27] = 0.12, *p* = .737), and the Outcome × Watch Period interaction was not significant (*F*[1.57, 42.39] = 0.37, *p* = .644). For the stimulus-locked, task-evoked pupil modulations there was a significant main effect of Outcome, *F*(1, 27) = 37.68, *p* < .001, *ηp^2^* = 0.58, but the main effect of Watch Period (*F*[2, 54] = 2.10, *p* = .132) and the Outcome × Watch Period interaction (*F*[2, 54] = 2.01, *p* = .144) were not significant. Simple main effects analysis showed that misses resulted in greater pupil modulation than correct rejections during Watch Period 1 (*F* = 30.69, *p* < .001) and Watch Period 2 (*F* = 8.02, *p* = .009), but not during Watch Period 3 (*F* = 1.20, *p* = 0.283).

**Figure 5.**
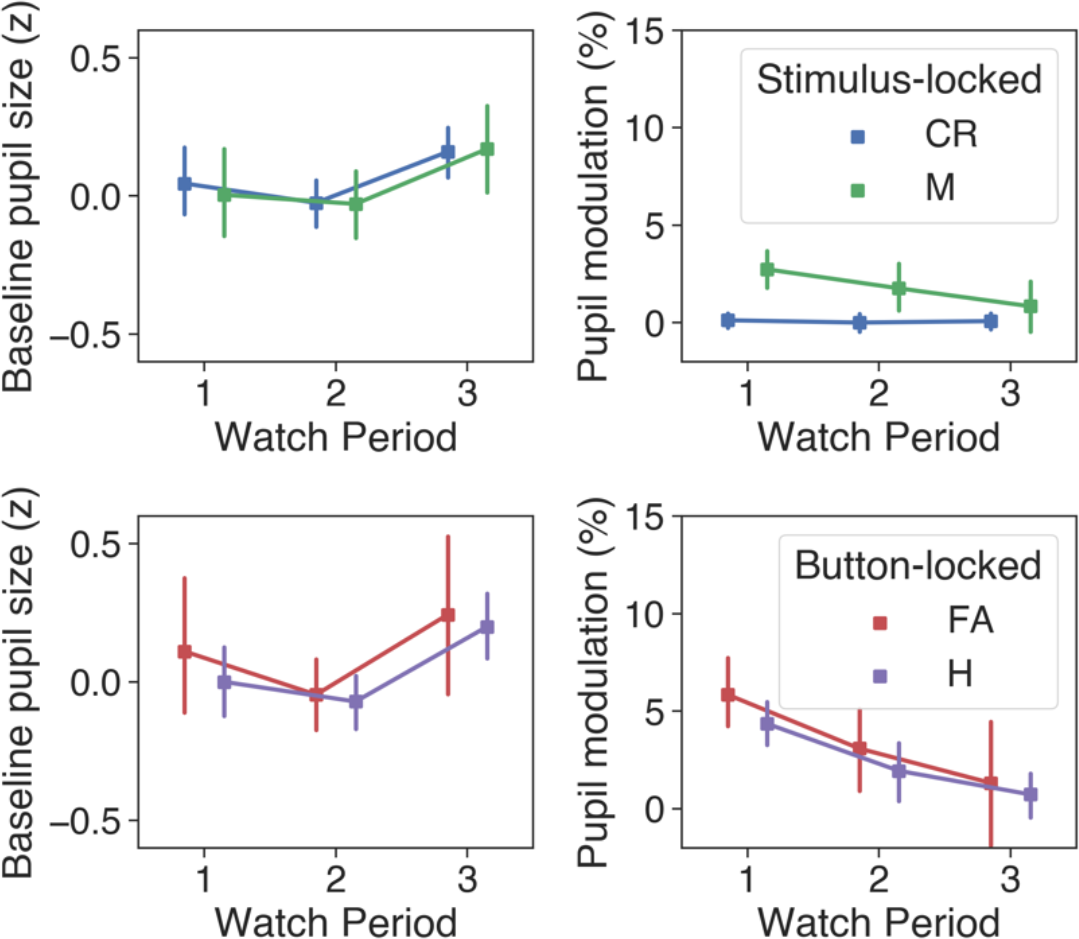
Stimulus- (M: misses; CR: correct rejections) and button-locked (H: hits; FA: false alarms) pupil averages across each Watch Period in Experiment 1. Error bars reflect 95% confidence intervals (bootstrapped, 1000 iterations).

For button-locked (i.e., hits and false alarms) baseline pupil data, the main effect of Watch Period was not significant, *F*(1.58, 33.25) = 2.32, *p* = 0.124, and the effect of Outcome was marginally significant, *F*(1, 21) = 4.03, *p* = .058, with baseline pupil size being greater on average for false alarms compared to hits. The Outcome × Watch Period interaction did not significantly effect baseline pupil size, *F*(2, 42) = 0.141, *p* = .869. For the button-locked pupil modulations, the main effect of Outcome was not significant, *F*(1, 21) = 0.004, *p* = .953, but there was a significant main effect of Watch Period *F*(2, 42) = 8.86, *p* < .001, *ηp^2^* = 0.3. Post hoc *t*-tests revealed that the average %-modulation for Watch Period 1 was greater than it was for Watch Period 2 (*MD* = 2.32, *t*[22] = 2.95, *p* = .023) or Watch Period 3 (*MD* = 4.04, *t*[22] = 3.74, *p* = .004), but that Watch Period 2 and Watch Period 3 did not differ significantly (*p* = .297). The Outcome × Watch Period interaction for button-locked pupil modulations was not significant (*F*[1.52, 31.87] = 0.09, *p* = .861).

Overall, these patterns in the pupil data are consistent with the prediction that the magnitude of task-evoked responses will mirror behavioral performance and decline as time-on-task increased.

#### 2.2.3 Correlational analyses

Across all button responses included in the analysis, RT did not significantly correlate with prestimulus baseline pupil size, *r*(4102) = -.009, *p* = .559, or task evoked pupil size, *r*(4102) = -.021, *p* = .073. In line with previous literature (e.g., see de Gee et al., 2014), there was a significant negative correlation between baseline and task evoked pupil size *r*(4102) = -.273, *p* < .001.

### 2.3 Discussion

This experiment examined the relationship between task performance and pupil size in a prolonged, uninterrupted vigilance task with visually presented stimuli. Participants monitored four centrally located equiluminant gratings continuously for 30 min under the instruction to respond by pressing the space bar every time they detected a target. The task required successive discrimination (Parasuraman, 1979), but the high background event rate, the brief target duration, and the temporal and spatial uncertainty of the targets added elements of difficulty (Broadbent, 1958; Mackie, 1987; Warm et al., 2008). In line with robust trends in the literature, we predicted that task performance and the magnitude of task-evoked pupillary responses would decrease as time-on-task increased.

Key behavioral measures were indicative of a classic vigilance decrement. Both the percentage of hits and the RT for hits changed across each successive 10 min watch period in a manner reflecting declining vigilance. These findings are consistent with well-established findings in the literature regarding the effects of time-on-task on detection performance under conditions of prolonged monitoring (e.g., Broadbent, 1953; Broadbent & Gregory, 1965; Buck, 1966; Mackworth, 1948, 1950, Parasuraman & Davies, 1976, 1982; Warm et al., 2008). The signal detection measures, sensitivity (*d’*) and response bias (*c*), were calculated to gain further insight into the cause of the declining percentage of hits. Given that the nature of the task was in making trivially easy judgements about suprathreshold stimuli, it is not surprising that sensitivity remained at ceiling throughout. However, there was a conservative shift in response bias, suggesting that the decline in accuracy was linked to the participants becoming less willing to report a detection, rather than a diminishing ability to discriminate targets from nontargets (Green & Swets, 1974). This is consistent with previous reports that the vigilance decrement in tasks with high event rates is more closely related to changes in the strictness of the decision criterion over time, rather than perceptual sensitivity (e.g., Baddeley & Colquhoun, 1969; Broadbent, 1971; Colquhoun, 1961; Parasuraman & Davies, 1976).

Event-related pupil data were extracted for all signal detection outcomes in order to gain insight into the cognitive processing associated with these events. For misses and correct rejections, data were time-locked to the onset of the relevant stimulus event (bottom-left panel of Figure 4). Notably, misses resulted in reliably greater pupil dilation than correct rejections, a similar observation to that made by Beatty (1982) in his auditory vigilance experiment. As Beatty (1982) suggested, from a signal detection perspective, an enhanced pupillometric response to missed targets may reflect increased processing of sensory information for stimuli that fall close to the decision criterion. In this vein, the pupil dilation following missed targets may in part reflect subconscious processing of the target stimuli (Laeng et al., 2012). However, in the context of this experiment, such an interpretation must be tempered against the possibility that pupil modulation for misses was linked to the detection, decision and motor effects of neighboring button-presses falling just outside of the permissible hit range. Previous research indicates that the pupillometric effects associated with motor acts can begin to emerge up to 1000 ms prior to the act itself (e.g., Einhäuser et al., 2010; Hupé et al., 2009; Martin et al., 2020; Richer & Beatty, 1985), which means that false alarms whose RT fell just outside the permissible hit range (upper bound of 1156 ms) may have contributed to the pupil dilation for missed targets. For hits and false alarms, pupil data were time-locked to the button response and showed typical patterns of modulation associated with motor preparation and execution. Pupil dilation was also significantly greater for false alarms compared to hits (lower-right panel of Figure 4). As suggested by Murphy et al. (2011), who had similar findings, this difference may reflect the cognitive effects of a self-regulatory performance monitoring process.

However, the larger pupil dilation for false alarms may also be associated with the higher degree of uncertainty that accompanies these events compared to correct detections (Yu & Dayan, 2005).

To examine the effects of time-on-task on pupil dynamics, scalar values of baseline and task-evoked pupil size were calculated for all stimulus and button events in each third of the task (Figure 5). As with Beatty (1982), baseline pupil size for all outcomes was relatively unchanged across the duration of the task, suggesting that the mode of organismic activation linked to fluctuations in tonic pupil diameter was not related to the central processes underpinning the vigilance decrement. In contrast to this, however, task-evoked pupil modulations for misses, hits and false alarms exhibited a marked decline in magnitude across each successive Watch Period. This pattern of change parallels the decline in vigilance indexed by the percentage of hits and RT for hits, and is therefore consistent with the findings from the vigilance experiments of Beatty (1982) and Murphy et al. (2011), as well as various other pupillometric studies of tasks requiring sustained attention (e.g., Hopstaken, van der Linden, et al., 2015; McIntire et al., 2014; Unsworth & Robison, 2016).

Task-evoked pupillary responses have been linked to phasic activation of the LC-NA system by neurophysiological and behavioral studies in both human and non-human primates (e.g., Alnaes et al., 2014; Aston-Jones & Cohen, 2005; Beatty, 1982; de Gee et al., 2017; Einhäuser, Stout, Koch, & Carter, 2008; Gilzenrat et al., 2010; Jepma & Nieuwenhuis, 2011; Joshi, Li, Kalwani, & Gold, 2016; Murphy, O’Connell, O’Sullivan, Robertson, & Balsters, 2014; Murphy et al., 2011; Phillips, Szabadi, & Bradshaw, 2000; Rajkowski, Kubiak, & Aston-Jones, 1993; Thorne et al., 2005; Urai, Braun, & Donner, 2017; Varazzani, San-Galli, Gilardeau, & Bouret, 2015). Further, single-unit recording studies in animals have found that phasic activation of the LC-NA system occurs typically in response to task-related events during periods of high performance (e.g., Aston-Jones, Chiang, & Alexinsky, 1991; Aston-Jones, Rajkowski, Kubiak, & Alexinsky, 1994; Rajkowski, Kubiak, & Aston-Jones, 1994). For example, the study by Aston-Jones et al. (1994) revealed that noradrenergic neurons in monkey LC are phasically activated by infrequent target cues during a vigilance task, and also that the amplitude of these phasic responses diminishes over time. With respect to the well-known functional association between task-evoked pupil responses and phasic LC activation (Laeng et al., 2012), the findings from these animal studies fit well with those from the present study.

From a theoretical standpoint, it is difficult to ascertain whether the findings of this study fit best with a resource depletion, mind wandering, or resource control-failure account of the vigilance decrement (see Caggiano & Parasuraman, 2004; Smallwood & Schooler, 2006; Thomson, Besner, & Smilek, 2015). Key indicators would be whether participants experienced the task as being effortful and the extent to which they engaged in task-unrelated thought, but we did not obtain these data as it would have required the use of intermittent thought probes (e.g., Hopstaken et al., 2015; Smallwood et al., 2004; Unsworth & Robison, 2016), which involve temporary disengagement and therefore undermine a key aspect of classical vigilance task design—the requirement for continuous monitoring (Parasuraman & Davies, 1976). Thought probes may also serve as ‘mini breaks’, which can disrupt task monotony and alleviate the vigilance decrement (Ariga & Lleras 2011; Ralph et al., 2016; Ross et al., 2014). In the absence of subjective reports, only the pupil data and the nature of the task can serve as a basis for inferring the cause of the vigilance decrement. First, the task itself was prolonged and monotonous, and participants were without a strong incentive to maintain high levels of vigilance—conditions that provide fertile grounds for mind wandering (Smallwood & Schooler, 2006). Second, the magnitude of task-evoked dilations reduced across the course of the task, an effect that has previously been linked to disengagement (Hopstaken, van der Linden, et al., 2015) and mind-wandering (Smallwood et al., 2011). Taken together, this suggests that the vigilance decrement in the current experiment may have been linked primarily to mind-wandering, or a resource control-failure leading to thought intrusion, but further data would be required to confirm this.

In sum, the present study replicated the well-known vigilance decrement—the reduction in detection performance that takes place during conditions of prolonged and continuous monitoring. Mirroring this behavioral effect, task evoked pupil responses declined across the duration of the task, but baseline pupil size was mostly unchanged, suggesting that the vigilance decrement may have been linked to gradual disengagement of attention as the task progressed, rather than a change in organismic arousal state.

## 3 Experiment 2

Traditional experimental vigilance tasks aim to emulate the conditions of real-world operator settings, but another form of vigilance task—the Psychomotor vigilance task (PVT: Wilkinson & Houghton, 1982)—aims to quickly assess declines in vigilant attention associated with sleep loss, circadian factors and other environmental stressors (Basner & Dinges, 2011; Basner, Mollicone, & Dinges, 2011; Blatter et al., 2006; Caldwell, Prazinko, & Caldwell, 2003; Dinges et al., 1997; Van Dongen & Dinges, 2005; Graw et al., 2004). In a PVT, instead of responding to infrequent signals over a prolonged period, subjects must make speeded responses to more regular signals occurring at pseudorandom intervals over a short period of time, usually 10 min or less. Wilkinson and Houghton’s (1982) original version of this task was administered on a small hand-held battery-powered device displaying a millisecond counter set to ‘000’. The subject held the device and quickly pressed a button every time the counter began to increment, which happened at intervals ranging between 1-10 s. Upon detection of a response, the timer froze for 1.5 s, and the RT was saved before the timer reset to ‘000’. A variety of performance metrics can be derived from the data produced by this task, but analysis commonly focuses on mean and median RT, the fastest and slowest 10% of trials, and the proportion of ‘lapses’, which are usually defined as RTs greater than 500 ms (Basner & Dinges, 2011).

In Experiment 2 we sought to examine how pupil measures relate to PVT task performance, but with a novel stimulus approach optimized for pupillometry. Most PVTs utilize the prototypical stimulus of a running millisecond timer that counts up from zero, but as noted by Thorne et al. (2005), this may have undesirable consequences. From a behavioral and pupillometric perspective, the two most relevant points made by Thorne et al. are as follows. First, the intensity of the stimulus changes in a nonlinear fashion as the running counter increases, which could be a source of increasing variance in both behavioral and pupil measures. Second, the counter provides feedback to participants whether feedback is desired or not, which may enable them to monitor their own performance during the session and increase attention or effort to compensate for a noticed decline—a boon that would typically not be available in real-world settings. To avoid these issues, Thorne et al. (2005) devised a version of the PVT with a luminance-based graphic stimulus comprising two alternating black and white circular annuli resembling a target or “bull’s eye”. This PVT was administered on a dedicated hand-held device and produced highly comparable results despite its different stimulus characteristics.

Since a luminance-based stimulus such as that used by Thorne et al. (2005) would trigger the pupillary light reflex, the RT-initiating stimulus opted for in the present PVT experiment was a change in the orientation of a low-contrast grating. Participants were instructed to monitor a grating at the center of a screen and respond as quickly as possible by pressing space whenever it flipped on its side (i.e., when it rotated 90°). As in other PVTs, the vigilance element of the task was instantiated with time-on-task (∼13 min) and ISI (4-12 s) parameters. Due to the use of a low intensity stimulus, we expected that RTs in the current PVT would be slower on average than for PVTs using a running counter stimulus. However, because our approach avoids the confounds of variable stimulus intensity and feedback, behavioral and pupil measures should more faithfully reflect changes in vigilant attention. Based on the general findings outlined in the introduction we predicted that time-on-task, both within and between successive blocks of the PVT, would lead to declining performance and a decrease in pupil size. Also, following the findings of Kristjansson, Stern, Brown, and Rohrbaugh (2009) and Unsworth and Robison (2016), who employed similar tasks, we expected that poorest performance would be associated with smaller pupils at baseline.

### 3.1 Materials and methods

#### 3.1.1 Participants

Twenty-five participants (18 females; age range 18-36 years, *M* = 22.96, *SD* = 4.65) completed the experiment voluntarily or in exchange for course credit. All participants were students at Swansea University reporting normal or corrected-to-normal acuity and color vision. The experimental protocol was approved by the Department of Psychology Ethics Committee at Swansea University and the Ministry of Defence Research Ethics Committee. Written informed consent was obtained from each participant.

#### 3.1.2 Design

Performance and pupil measures were analyzed in a repeated measures design as a function of Trial Group (1, 2, 3, 4, 5: sets of 18 contiguous trials within a PVT block) and Block (1, 2, 3: successive blocks of 90 trials)—two factors whose purpose was to capture within- and between-block effects of time-on-task. The ISI, defined as the period between the last button response and the next flip of the grating, varied between 4-12 s. This period included a fixed component of 2000 ms and a random component varying between 2-10 s. The random component was constrained such that a third of the intervals would be short (2000-4666 ms), a third medium (4666-7333 ms), and a third long (7333-10000 ms). The grating was present at the center of the screen throughout the task.

#### 3.1.3 Stimuli and apparatus

The stimulus was a grating enveloped within a cosine window (spatial frequency = 0.1, *SD* = 12, 39% contrast) at the center of the screen, spanning 2° × 2° of visual angle. It was generated using the same process as specified for Experiment 1. All other details relating to hardware and software were the same as for Experiment 1.

#### 3.1.4 Procedure

Participants completed 3 consecutive blocks of the PVT in a single testing session, taking a forced break of only 1 min between blocks. In each block, participants were instructed to monitor the ‘circular stimulus’ at the center of the screen and to respond as quickly as possible by pressing the space bar whenever it flipped on its side, thereby to reset it to its original position (see Figure 6). Each block lasted approximately 13 min (*M* = 12.8, *SD* = 0.63), with some small variability arising from differences in RT and the random element of the ISI. Continuous recordings of gaze position and pupil data were obtained for each trial, and RT was defined as the time in milliseconds between the flip of the grating and the subsequent button response. A 5-point calibration and validation routine was performed at the start of each block. The task was performed in a dimly lit room.

**Figure 6.**
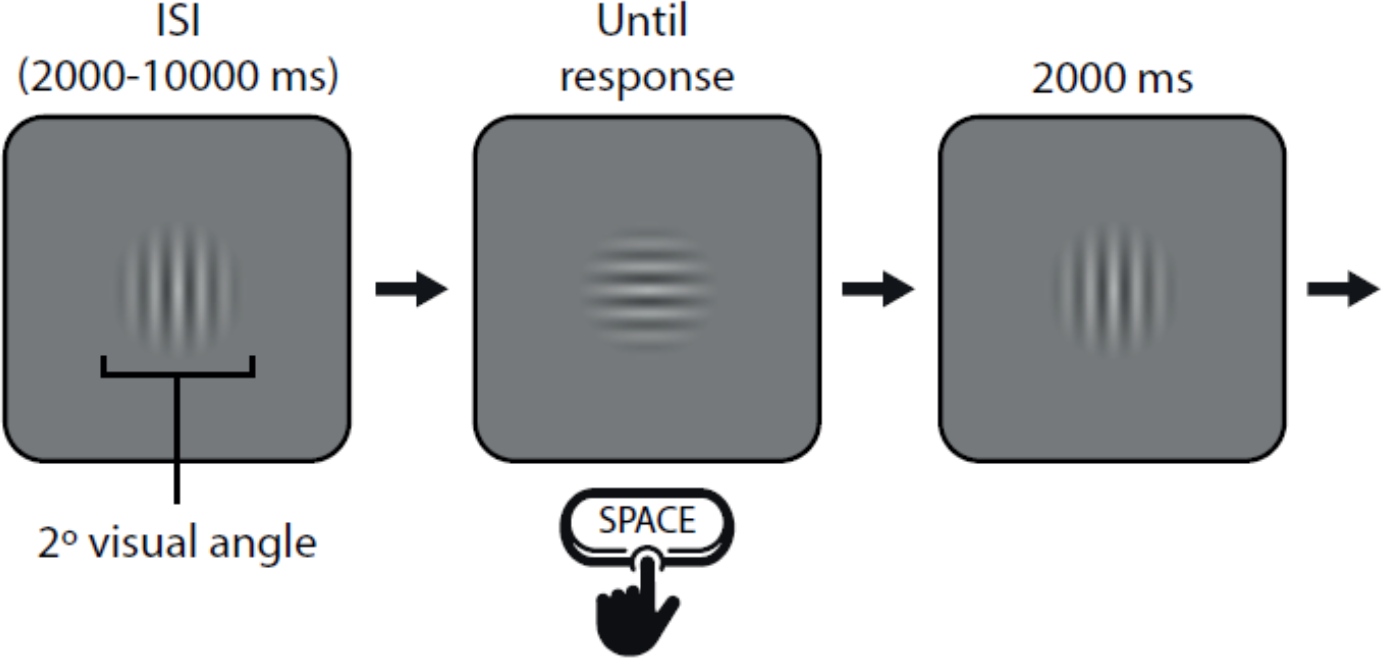
Stimuli and trial sequence for Experiment 2. Participants monitored a grating and responded by pressing space every time it flipped on its side (every 4-12 s).

#### 3.1.5 Data processing and statistical analysis

Pupil data were processed using the same general approach as described for Experiment 1. The average amount of data replaced by blink interpolation across all participants included in the analysis was 6.82%. Segments of pupil data 2500 ms in length were extracted for each trial, time-locked to the button response (-1000 to 1500 ms). These data were expressed as %-modulation from a baseline defined as the average pupil size in a 500 ms period prior to the RT-initiating stimulus event.

To assess task performance we focused on 1/RT and lapse frequency, which are among the most sensitive measures of alertness in PVTs (Basner & Dinges, 2011). Lapses in PVTs are traditionally defined as RT greater than 500 ms but due to our novel take on the task we defined lapses as RT greater than two median absolute deviations (Leys et al., 2013) from each participant median, which resulted in an average lapse threshold of 585 ms (*SD* = 109) across participants. Our pupil measures of interest were baseline pupil size and the task-evoked pupil response. Baseline pupil size was defined as the average *z*- transform of pupil size in the baseline period, whereas the task-evoked pupil response was defined as the average percentage of pupil modulation around the time of the button response (-500 to 1500 ms). The data for each of these four variables were analyzed separately using two-factor (Trial Group × Block) repeated measures ANOVA. Where Mauchly’s *W* indicated that the assumption of sphericity was violated, *p*-values were adjusted using the Greenhouse-Geisser correction. We conducted further analyses on pupil measures for the fastest and slowest 20% RTs to determine whether pupil size at baseline was indicative of faster or slower detection responses (Kristjansson et al., 2009; Unsworth & Robison, 2016), and more generally how the extremes of performance are reflected in the pupil data. The mean and standard deviation of horizontal (*M* = 518, *SD* = 10) and vertical (*M* = 388, *SD* = 14) gaze position for all samples included in the analysis indicate that participants maintained steady fixation at the center of the screen throughout the task.

#### 3.1.6 Exclusions

Two participants were excluded from the analysis for yielding poor quality pupil data (both had over 50% interpolated data for baselines and over 60% interpolated data for task-evoked responses). The general pattern of results was the same both with and without the exclusion of these participants. For the pupil analyses, trials were excluded if there was a blink in the baseline or if more than 25% of data were interpolated across the whole epoch (28.06% of trials).

### 3.2 Results

#### 3.2.1 Task performance

Average RT across all participants was 420 ms (*SD* = 79) for non-lapse trials and 960 ms (*SD* = 1463) for lapse trials. ANOVA on mean 1/RT (i.e., the reciprocal transform of RT) revealed a significant Trial Group × Block interaction, *F*(8, 176) = 3.16, *p* = .002, *ηp^2^* = 0.13, the nature of which is illustrated in the top-left panel of Figure 7. Simple effects showed that 1/RT decreased significantly across Trial Group in each Block (Block 1: *F* = 16.65, *p* < .001; Block 2: *F* = 2.89, *p* = .027: Block 3: *F* = 3.40, *p* < .012), which is consistent with the prediction that performance would decline as time-on-task increased. Post hoc analysis with Bonferroni adjustment showed that, in Trial Groups 1-3, 1/RT was significantly greater in Block 1 compared to Blocks 2 and 3 (all *p*s < .05) and that 1/RT in Trial Group 4 was significantly greater for Block 1 compared to Block 3 (*p* < .05). No other comparisons were significant (*p* > .05). Therefore, as indexed by 1/RT, performance was best overall in Block 1 compared to Block 2 and Block 3, but the magnitude of this effect decreased across Trial Groups.

**Figure 7.**
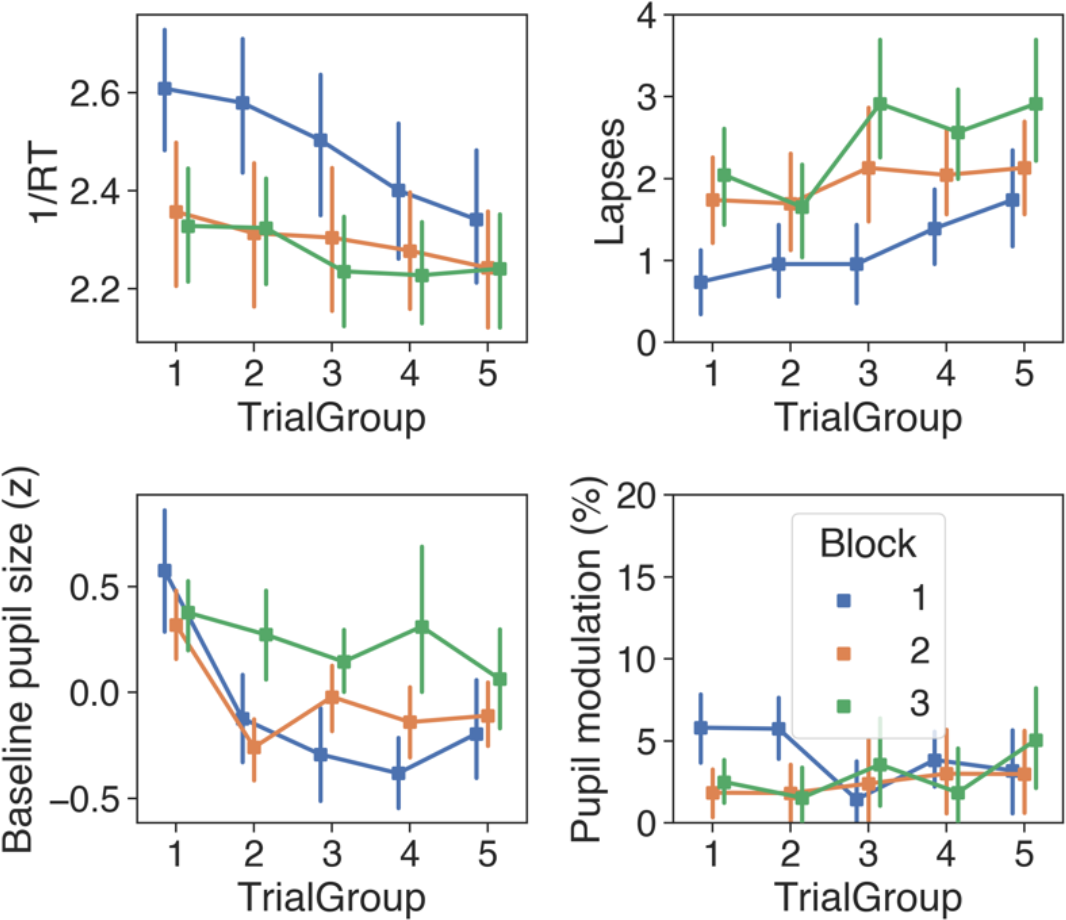
Performance (top row) and pupil (bottom row) measures across Trial Group and Block in Experiment 2, with error bars showing 95% confidence intervals (bootstrapped, 1000 iterations).

Average lapse frequency across participants was 27.6 (*SD* = 7.8). ANOVA showed that the main effect of lapse frequency was significant for Trial Group, *F*(4, 88) = 5.43, *p* < .001, *ηp^2^* = 0.20, and for Block, *F*(2, 44) = 16.74, *p* < .001, *ηp^2^* = 0.43 (top-right panel of Figure 7), but that the Trial Group × Block interaction was not significant (*F*[8, 176]*, p* = .541). Simple effects for Trial Group showed that the number of lapses increased significantly throughout Block 1 (*F* = 3.16, *p* = .018) and Block 3 (*F* = 3.01, *p* = .022), but not Block 2 (*F* = 0.60, *p* < .662). Lapse frequency therefore followed the same general pattern as 1/RT, and together these data are consistent with the prediction that performance would decline as time-on-task increased.

#### 3.2.2 Pupil data

Grand-average button-locked pupil traces for each Block are shown in Figure 8. The pupil began to dilate slowly following the stimulus event and then rapidly after the button-press. In the 1500 ms following the button-press there was an average modulation of 5.22% and a peak latency of 880 ms. A conspicuous trough in the pupil traces after the button-press coincides with a transient but marked increase in the percentage of interpolated data. This artifact resembles the blink-induced pupillary response (e.g., Knapen et al., 2016) and is therefore indicative of task-correlated blinking (i.e., participants tended to blink after button presses). We did not correct this artifact with linear interpolation as it would involve altering too much data and excluding more trials.

**Figure 8.**
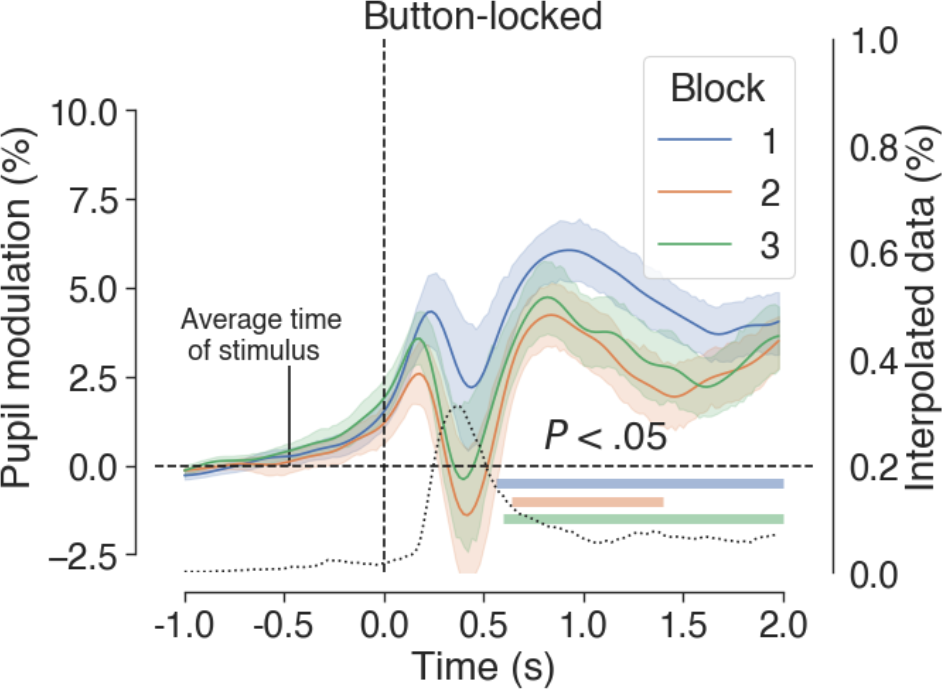
Average button-locked pupil traces for each Block in Experiment 2. The black dotted trace shows the percentage of interpolated data, which indicates task-correlated blinking. Shaded areas surrounding the colored traces show the SEM (bootstrapped, 5000 iterations) and colored horizontal bars denote clusters of significant modulation from baseline, as revealed by nonparametric permutation tests (1024 permutations, *p* < .05, cluster-corrected for multiple comparisons).

ANOVA on the baseline pupil measures revealed a significant Trial Group × Block interaction, *F*(3.81, 76.10) = 3.47, *p* = .013, *ηp^2^* = 0.15, which is displayed in the bottom-left panel of Figure 7. Simple effects analysis for Trial Group showed that baseline pupil size decreased significantly across Block 1 (*F* = 13.57, *p* < .001) and Block 2 (*F* = 7.02, *p* < .001), but not Block 3 (*F* = 0.89, *p* = .470). Bonferroni-corrected post hoc *t*-tests revealed that baseline pupil size was significantly greater in Trial Group 4 for Block 3 compared to Block 1 (*p* = .007), but no other between Block comparisons were significant (*p* > .05).

For measures of pupil modulation, there was a significant main effect of Block, *F*(2, 42) = 5.93, *p* = .010, *ηp^2^* = 0.23, but the effect of Trial Group (*F*[4, 80] = 1.86, *p* = .125) and the Trial Group × Block interaction (*F*[4.83, 96.63] = 1.87, *p* = .109) were not significant. Simple main effects showed that pupil modulation was greater in Block 1 for Trial Group 1 (*F* = 8.97, *p* < .001) and 2 (*F* = 9.07, *p* < .001), but not for Trial Groups 3 to 5 (*p* > .05).

#### 3.2.3 Correlational analyses

Across all trials included in the analysis, RT correlated significantly with task evoked pupil size, *r*(4466) = -.159, *p* < .001, but not with baseline pupil size, *r*(4466) = -.011, *p* = .459. The classic negative correlation between baseline and task evoked pupil size (e.g., see de Gee et al., 2014) was also present, *r*(4466) = -.366, *p* < .001.

3.2.4 – Fastest vs. slowest RTs

To explore how the extremes of performance are reflected in the pupil data we conducted further analysis on trials with the fastest (*M* = 347 ms, *SD* = 43 ms) and slowest (*M* = 746 ms, *SD* = 43 ms) 20% RTs. Figure 9 displays the pupillometry results for these extreme quintiles. Baseline pupil size did not differ significantly (*p* > .05, left panel of Figure 9), but there was a significant difference in the button-locked pupil traces (*p* < .05, cluster-corrected permutation test, right panel of Figure 9) marked by a cluster spanning the button event. This difference clearly pertained to the timing and magnitude of pupil dilation. For the slowest RTs, dilation began prior to the button response, whereas for the faster RTs dilation did not begin until afterwards. Further, the average modulation in the 2000 ms post-button period was greater on average for the slowest (*M* = 4.39%, *SD* = 5.46%) than for the fastest RTs (*M* = 2.45%, *SD* = 3.23%). These data do not corroborate previous reports of pretrial baseline predicting performance (e.g., Kristjansson et al., 2009; Unsworth & Robison, 2016), but rather they suggest that, at least within the context of our experiment, the pattern of pupil dilation prior to a detection response may be the more relevant predictor. In this respect, our data are in line with recent PVT studies where the fastest RTs were associated with larger pupil dilations in the ISI (Unsworth et al., 2020; Unsworth & Robison, 2018).

**Figure 9.**
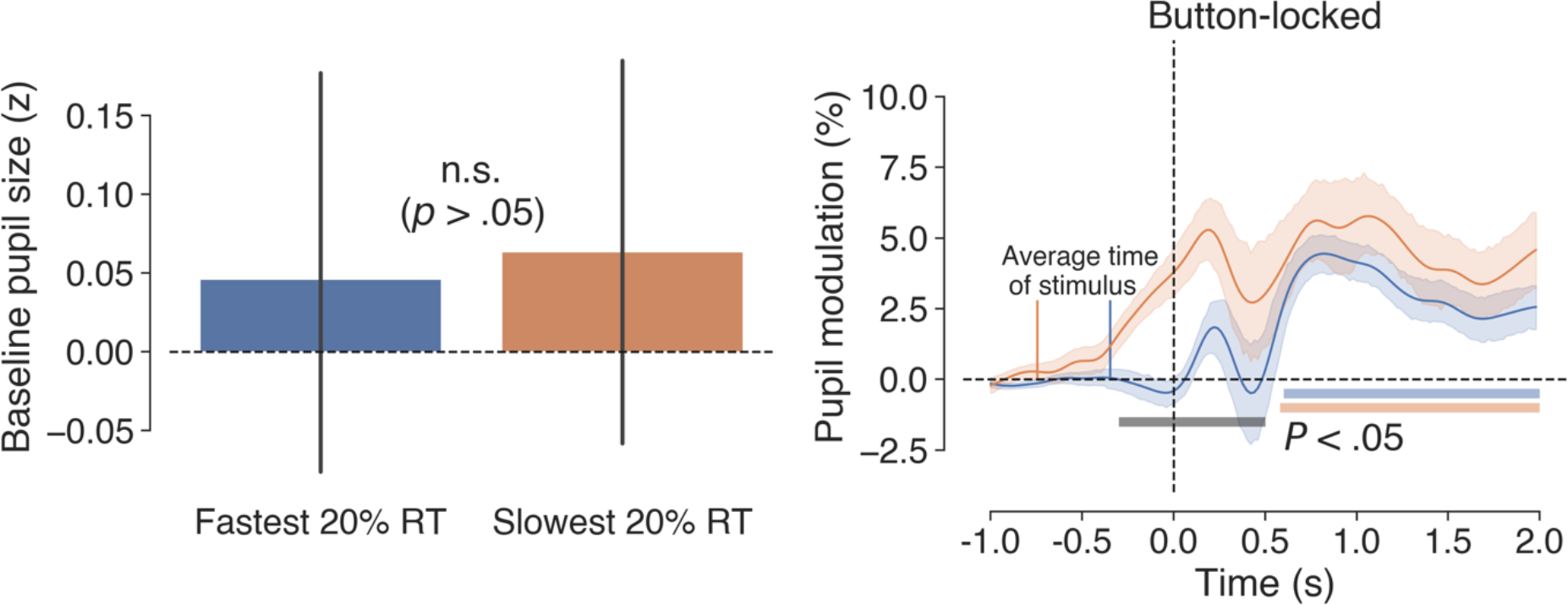
Baseline and button-locked pupil measures for the fastest (*M* = 347 ms, *SD* = 43 ms) and slowest (*M* = 746 ms, *SD* = 43 ms) 20% RTs in Experiment 2. The left panel shows mean prestimulus baseline pupil size with 95% confidence intervals (bootstrapped, 1000 iterations) and the right panel shows pupil dilations time locked to button responses, with shaded areas surrounding the pupil traces denoting the SEM (bootstrapped, 5000 iterations). Colored horizontal bars in the right-hand panel denote clusters of significant modulation from baseline for the respective traces (grey bar represents the difference between the traces), as revealed by nonparametric permutation tests (1024 permutations, *p* < .05, cluster-corrected for multiple comparisons).

### 3.3 Discussion

This experiment sought insight into the relationship between pupil size and performance measures in a novel PVT. Participants monitored a low contrast grating for a sudden 90° rotation and responded with a button press as quickly as possible after the event. We adopted an atypical stimulus approach to avoid confounds associated with the canonical running counter stimulus—namely its variable intensity and the performance feedback that it provides—which could potentially contribute to variance in behavioral and pupillometric measures (Thorne et al., 2005). Participants completed three successive blocks of the task taking only a 1-min break in between, and changes in performance and pupil measures were explored both within and between blocks. We predicted that performance and pupil size would decrease as time-on-task increased, and that worse performance would be associated with smaller pupils at baseline.

The initial point to note is that our novel stimulus approach led to longer RTs than are typically observed in PVTs that use the canonical running counter stimulus. In these tasks, average RT for subjectively alert participants is generally in the range of 200 to 300 ms (e.g., Basner, Mollicone, & Dinges, 2011; Blatter et al., 2006; Dorrian, Roach, Fletcher, & Dawson, 2007; Loh, Lamond, Dorrian, Roach, & Dawson, 2004; Matsangas, Shattuck, & Brown, 2016; McClelland, Pilcher, & Moore, 2010; Wilkinson & Houghton, 1982), whereas in the current PVT, also with subjectively alert participants, average RT was 420 ms. We attribute this to differences in stimulus intensity. The running counter stimulus is dynamic and constantly changing, providing a constantly refreshed cue for the participant to respond, whereas a change in the orientation of a low contrast grating is more subtle and discrete, and issues no refreshing cue to respond.

As predicted, the main performance measures exhibited typical time-on-task effects, with 1/RT decreasing and the number of lapses increasing as time-on-task increased. This general pattern was observed within and between each block of the PVT for both performance measures. For 1/RT, the biggest change was between the first block and the two subsequent blocks, with the difference being largest across the first three groups of trials. Lapse frequency increased gradually within each block and between successive blocks. These patterns in the performance data were statistically robust even without the state manipulations (e.g., time of day, sleep deprivation) and large number of repeated tests that are often integral to the design of mainstream PVT research (e.g., Basner et al., 2011; Blatter et al., 2006; Dorrian et al., 2003; Graw, Kräuchi, Knoblauch, Wirz-Justice, & Cajochen, 2004; Loh et al., 2004; Manousakis, Maccora, Ftouni, & Anderson, 2017).

As regards the pupil data, the pattern of within-block declining baseline pupil size broadly reflected the decline in task performance, corroborating findings from previous PVT studies (Massar et al., 2016; Unsworth & Robison 2016) as well as various other studies which examined pupil and performance measures in vigilance or sustained attention (van den Brink et al., 2016; Grandchamp et al., 2014; Hopstaken, van der Linden, et al., 2015; McIntire et al., 2014; Van Orden et al., 2000). However, the relationship between baseline pupil size and task performance was not clear cut. Participants performed best and had the largest baseline pupil size at the beginning of each block, but the sharp drop in baseline pupil size which occurred between the first and second Trial Group did not have a commensurate drop in performance. Further, baseline pupil size was largest overall and showed the least variability across Trial Groups in Block 3, where performance was at its worst. In a similar fashion, the task-evoked pupil responses were largest at the beginning of Block 1, where performance was best, but were less consistent with respect to the performance data at other times. These patterns in the pupil data are in line with the general prediction that pupil size would decrease as time-on-task increased, but they run contrary to the prediction that worse performance would be reflected in smaller pupils at baseline.

Previous experiments offer conflicting evidence as to whether optimal task performance is associated with larger or smaller pupils at baseline (e.g., Kristjansson et al., 2009; Unsworth & Robison, 2016). To address this issue, we compared baseline and task-evoked pupil responses for the trials with the fastest and slowest 20% RTs. Whilst there was no significant difference in baseline pupil size between these two groups of trials, there was a clear difference between the observed pupil traces. For the faster RTs, dilation did not begin until after the button response was made, whereas for the slower RTs, dilation was apparent around 500 ms before the button response and increased gradually until it peaked shortly afterwards. The finding of gradual dilation prior to an overt detection response, which is well documented in the literature, has been linked to cognitive factors associated with target recognition and decision making (e.g., see Einhäuser et al., 2010; Martin et al., 2020; Richer & Beatty, 1985). The reason we see this only for the slowest and not the fastest trials probably reflects the difference in RT and the fact that genuine cognitive effects on pupil size tend not to develop until at least 220 ms from the causal event (Mathôt et al., 2015, 2018). For the fastest trials, pupil modulation effects relating to target recognition and decision making were likely mixed in with the motor component.

The findings from the current experiment are generally consistent with previous studies showing time-on-task effects on performance and pupil size, but they do not align perfectly with a specific theory of vigilance. The understimulating and unrewarding nature of the task does however provide ripe conditions for mind-wandering, suggesting that this may have been partly responsible for the decline in performance. Previous studies have also reported larger pupils at baseline during periods of mind-wandering and poor task performance (e.g., Franklin, Broadway, Mrazek, Smallwood, & Schooler, 2013; Smallwood et al., 2011, 2012; Unsworth & Robison, 2016), which is the pattern that was observed in Block 3 of the current experiment. Research also suggests that very short breaks can reduce mind-wandering and lead to performance improvements by temporarily boosting motivation (e.g., Ariga & Lleras, 2011; Ralph, Onderwater, Thomson, & Smilek, 2016; Ross, Russell, & Helton, 2014), which fits with the pattern of data in the current experiment, where participants’ performance was restored to more-optimal levels after taking a 1-min break in between each block.

We recognize that various factors relating to the individual state of the participants could have influenced the results of the present experiment. For example, performance in PVTs is affected by sleep pressure (Blatter et al., 2006), time-of-day and its interaction with circadian rhythms (Van Dongen & Dinges, 2005; Graw et al., 2004), the consumption of stimulants such as caffeine (Van Dongen et al., 2001), and individual differences in intrinsic alertness (Unsworth et al., 2020). The current experiment did not control for any of such factors, but this could easily be achieved in a subsequent study. For instance, circadian effects could be controlled for by excluding strong “morning and evening types” (Horne, Brass, & Pettitt, 1980) and by testing participants at the same times during the day, after they have reported having similar amounts of sleep. Alternatively, one could examine how performance and pupillometry vary with respect to individual differences in a broad range of cognitive and self-reported personality factors (e.g., Unsworth et al., 2019, 2020).

Finally, we note that our novel take on the PVT limits the extent to which it can be directly compared to a more traditional PVT. The use of an alternative stimulus was desirable to avoid certain confounds, but the experiment also differed in terms of block length and ISI. In their general recommendations for the standardized design and analysis of PVTs, Basner and Dinges (2011) suggest using an ISI of 2-10 s and having a fixed block length of 10 min. Due to the way the current experiment was implemented, ISI was 4-12 seconds and block length was variable (*M* = 12.8 min, *SD* = 0.63 min). Future experiments may wish to bring our approach closer to the task specifications set out by Basner and Dinges (2011), which would broaden the basis for comparison of experimental findings in the wider literature.

## 4 General discussion

Recent pupillometric studies of vigilance and sustained attention suggest that measurements of pupil size could potentially be used in operational settings to monitor performance, and perhaps even to predict and prevent errors associated with lapses of attention before they occur. But the literature in this area—especially regarding visual tasks—is sparse, and differences in methodology and task requirements have led to conflicting findings. The purpose of the current study was to further explore the relationship between pupil size and performance measures in the context of well-established task frameworks from the vigilance literature.

The most consistent finding across both experiments regarding the relationship between pupil size and monitoring performance was that, in line with previous experimental findings (e.g., Beatty, 1982; Hopstaken et al., 2015; Unsworth & Robison, 2016) and the predictions of established theory (Aston-Jones & Cohen, 2005), task-evoked pupil responses were generally more pronounced when performance was best. This trend was most consistent in Experiment 1, where the decline in detection performance was mirrored by a decline in the magnitude of task-evoked responses associated with hits, misses, and false alarms. In Experiment 2, the relationship between task-evoked responses and performance measures was less consistent, although the largest responses did occur when performance was best (i.e., at the beginning of Block 1). In general, these findings suggest that changes in task-evoked pupil responses may serve as an accurate indication of general task engagement, with a decline in their magnitude over time reflecting cognitive disengagement from the task and an increased likelihood of suboptimal performance.

Our baseline pupil measures did not show a consistent relationship with performance. In Experiment 1, baseline pupil size was mostly unchanged across three successive periods of watch, despite a marked decrement in performance. In Experiment 2, baseline pupil size showed an overall decline within each Block, although the slope became less pronounced with each successive Block. Interestingly, baseline pupil size was biggest overall at the beginning of each Block, where task performance was best, suggesting that it reflects heightened arousal, alertness, and focused attention. But, by this account, our baseline measures in the PVT reflect combinations of autonomic tone as well as task-related factors, which means that they are not serving uniquely as a window of insight into the “tonic” mode of LC activation, as is often explicitly or implicitly assumed (see below). The lack of consistency in our baseline measures and their relationship with performance metrics is not unprecedented in light of the literature reviewed in the introduction, which indicates that the relationship is complex and in need of further characterization. One possibility raised by van den Brink et al. (2016) is that the effects of time-on-task on baseline pupil size obscure a more nuanced relationship with performance. In their gradual-onset performance task, after regressing out the effects of time-on-task from the baseline pupil data, the authors observed a quadratic relationship with performance, such that performance was optimal when baseline pupil size was at intermediate levels. This idea dovetails with the Yerkes-Dodson law (Yerkes & Dodson, 1908) of optimum arousal, whereby the relationship between task performance and arousal is described by an inverted-U function, such that poor performance is associated with both under- and over-arousal, and optimum performance occurs at a “sweet spot” on the arousal curve.

We refrained from using the words “tonic” and “phasic” to describe our pupil measures because we are aware of numerous caveats to the assumption that baseline and task-evoked measures map neatly onto the different modes of LC output. Joshi and Gold (2020) discuss this issue in detail and emphasize that, in the context of LC activation, the terms “tonic” and “phasic” differentiate between distinct modes of activation, and not simply between baseline and transient activity (Aston-Jones & Cohen, 2005).

Further, the operational definition of “tonic” and “phasic” pupil measures varies substantially between publications. Also, the precise neural mechanisms of the relationship between pupil measures and LC activation are presently unclear and it is possible that a third variable, as of yet not understood, may account for the observed pupil-LC link (Costa & Rudebeck, 2016).

In conclusion, the results of our two vigilance experiments support the general notion that changes in task-evoked pupil measures can be used to gain insight into monitoring performance in long and demanding tasks where the emphasis is on additive effects over a series of trials. But there is clearly a need for further research to determine the practical feasibility of utilizing pupil size as a psychophysiological marker of attentional lapses in real-time monitoring systems. Characterizing the precise relationship between different measures of behavioral performance, task-related factors and patterns of pupil behavior will be a crucial next step in this regard.

## Acknowledgements

The research was funded by a grant from the Defence Science and Technology Laboratory (DSTLX1000083208)—an executive government agency that ensures innovative science and technology contribute to the defence and security of the UK. We thank three anonymous reviewers for their comments and suggestions.

---

**Figure X.**
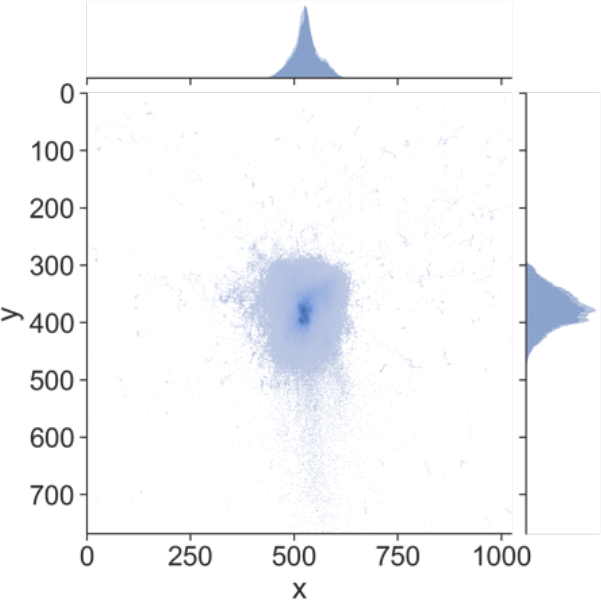
Joint histogram for x and y of gaze position in screen pixel coordinates.

## Footnotes

1 The distinction between successive and simultaneous discrimination tasks was first made by Parasuraman (1979). Successive tasks are absolute judgement tasks where observers must compare the current sensory input with a template in working memory in order to determine whether a particular stimulus is, or is not, a critical signal. Simultaneous tasks on the other hand are comparative judgement tasks, where each stimulus contains all of the information required to determine whether it is (or is not) a signal. Due to the involvement of working memory, successive tasks are thought to be more resource demanding than simultaneous tasks.

2 https://www.cogsci.nl/gabor-generator

## Notes

https://osf.io/yujw6/?view_only=902f6e591fa34640b2ea12c479f1940b

### Competing Interest Statement

The authors have declared no competing interest.

